# A simple technique to create pre-vascularised ‘delivery scaffolds’ for organoid therapies

**DOI:** 10.1101/2025.11.05.686546

**Authors:** Kevin X Cao, Nazanin Owji, Amna Kaizer, Gideon Pomeranz, Karen Price, Navroop Johal, David A Long

**Affiliations:** UCL Great Ormond Street Institute of Child Health

## Abstract

The growth in complex cellular constructs such as organoids for disease-modelling or regenerative therapies creates a need to overcome the problem of nutrition-delivery and surgical delivery. This logically requires a ‘pre-vascularised scaffold’, something that can embed and sustain the growth of these constructs and minimise the time needed for blood supply integration on implantation.

We present here a series of experiments and their techniques that demonstrate a low-cost fabrication pipeline to create biologically-compatible scaffolds alongside evaluations of their optimal parameters in-vivo, on-CAM and in-vitro using mice and the chick chorioallantoic membrane (CAM) assay as a rapid throughput testbed of scaffolds.

Utilising polylactic-glycolic acid (PLGA) scaffolds, we found an optimal range of porosity (300-500um, variable pore size), optimal lactide to glycolide ratio (82:18) and an approximation of the optimal cell seeding density using reset-vascular endothelial cells (R-VECs). Finally, we demonstrate a method to delineate the internal vascular network of these scaffolds with micro-computer-tomography (MicroCT), a low-cost analytical method that may become useful for network analysis in regenerative medicine research.

## INTRODUCTION

The growth of complex cellular, tissue, and organoid-based research for disease modelling and regenerative therapies has created a demand for simple methods to fabricate pre-vascularised scaffolds (PVS). Pre-vascularised scaffolds (PVS) are crucial for supporting advanced cellular constructs, facilitating nutrient delivery during in vitro culture, and ensuring rapid vascular integration (inosculation) upon implantation. Organoids face two key challenges in this context. First, they require a robust vascular network to deliver nutrients during in vitro growth and enable rapid inosculation upon implantation, allowing host vessels to connect without penetrating deeply into the construct. Second, a suitable transport platform is necessary to prevent structural damage during handling, ensuring safe delivery even under surgical conditions.

Vascular networks develop most effectively in interconnected channels between 200–400 µm in diameter (Laschke, Vollmar & Menger, 2009). Constructing these networks need not be costly or complex; PVS can be fabricated from low-cost polymer materials, with or without connective tissue components like collagen or decellularised matrices. Regardless of fabrication method, an ideal scaffold should maintain its form factor and promote neo-angiogenesis.

This study investigates scaffold porosity, in vivo structural integrity, and the use of Reset Vascular Endothelial Cells (R-VECs) for creating pre-vascularised scaffolds (Palikuqi et al., 2020). We also explore embedding blood vessel organoids into PVS and present a novel imaging method for visualising vascular networks in chick chorioallantoic membrane (CAM) assays using micro-computed tomography. Importantly, these methodologies are designed to be low-cost and accessible, offering a “recipe” for researchers to create PVS without the need for expensive cell culture platforms.

We aim to provide researchers with simple, cost-effective tools to create and evaluate PVS, enabling progress in regenerative medicine and tissue engineering.

## METHODS

### Fabrication of Polymer Scaffolds

Porous polymer scaffolds were fabricated using porogen-leaching or electrospinning, combining well-established techniques to produce scaffolds with defined porosity or fibrous matrices resembling the extracellular environment. PLGA was selected for its structural integrity and safe, non-toxic degradation in vivo, with L:G ratios of 75:25 and 82:18 tested at a concentration of 10% w/v.

### Porogen-Leaching Method

For porogen-leaching, PLGA was dissolved in 1,4-dioxane (or HFIP when hybridising with collagen) at 50°C for 1 hour. Spherical porogens (200–600 µm) were evenly distributed in a PTFE dish, ensuring consistent pore size and connectivity, while avoiding suboptimal vertex-to-vertex or edge connections common with salt porogens (Zhang et al. 2005). The PLGA solution was poured over the porogen layer, frozen at -80°C, and freeze-dried overnight (-1 atm, -80°C). Porogens were leached in warm water (37°C) with magnetic stirring for 2–4 hours, yielding porous scaffolds, which were cut into circular discs (1 cm diameter, 2–3 mm depth). Gelatin porogens required higher temperatures (60°C) and longer dissolution times, while salt porogens dissolved more rapidly but resulted in less optimal pore connectivity.

### Electrospinning Method

Fibrous scaffolds were created via electrospinning. PLGA was dissolved in an 80:20 dichloromethane (DCM) and N,N-dimethylformamide (DMF) solvent mix (10% w/v) and homogenised with magnetic stirring. The polymer solution was electrospun using a 16G nozzle under optimised conditions: a flow rate of 85 ml/min, applied voltage of 20 kV, and a firing distance of 20 cm. Environmental conditions were controlled at 23°C and 50% humidity. The resultant fibre sheets were cut into discs for use as scaffolds.

### Collagen Hybridisation

To enhance biological relevance, scaffolds were hybridised with collagen. A 4:1 collagen blend (10% w/v in HFIP) was employed, offering superior integration compared to dip-coating methods, likely due to (Sadeghi-Avalshahr et al. 2017).

### CAM-mounting, perfusion studies on-CAM

The chick chorioallantoic membrane (CAM) assay was used as a rapid, highly-angiogenic, cost-effective testbed for scaffold vascularisation and biocompatibility. This approach integrates established CAM techniques to enable high-throughput scaffold evaluation. Fertile chicken eggs (Henry Stewart & Co. Ltd) were cleaned with 50% ethanol and incubated at 37.5°C (Marsh Automatic Incubator). On day 3, embryos were examined for viability and prepared for scaffold implantation.

#### Preparation of CAMs

On day 3, 1 ml of egg albumin was withdrawn using a 14G syringe to lift the embryo closer to the eggshell. A 2.5 × 2.5 cm window was made in the shell using a blade and sealed with adhesive tape. Eggs were returned to the incubator. An alternative ex-ovo method incubated CAMs in jars on low-density polyethylene (LDPE), though this had higher mortality during scaffold mounting.

#### Scaffold Application and Harvesting

On day 10, scaffolds were placed at a small vascular injury site on peripheral CAM vessels. To minimise infection, 500 µL of phosphate-buffered saline (PBS) with 1% penicillin-streptomycin was applied. A 1 cm silicone ring was used to stabilise the CAM.

After 24 hours (day 11), scaffolds were harvested with a surrounding albumin rim and preserved in 4% paraformaldehyde (PFA). For some analyses, injections were performed prior to harvesting.

### Vascular mapping

Images were taken with a stereomicroscope of harvested scaffolds. ImageJ software was used to calculate the percentage coverage area of blood vessels after first converting RGB images to black and white binary images. Particle analysis function was used to calculate the percentage of pixels representing vascular networks as a percentage of all pixels.

### Micro-CT Imaging and Analysis

Three-dimensional imaging of tissue constructs was performed using an optimised in-ovo perfusion method with the radio-opaque contrast agent MicroFIL, followed by micro-computed tomography (Micro-CT) imaging (Woloszyk, Liccardo & Mitsiadis 2016). Traditional clearing techniques with wholemount immunostaining were unsuitable due to the inability of laser microscopy to penetrate polymer-based structures at depth. This method enabled effective visualisation and analysis of vascularisation within the scaffolds.

### Mouse Skin-Fold Scaffold Implantation Surgery

Ten-week-old male CD1 mice (Charles River) were used to evaluate all three PLGA scaffold compositions. Mice were anaesthetised with isoflurane, and dorsal hair was shaved and cleaned with chlorhexidine. Two scaffolds were implanted per mouse, one into each dorsal skin flank, via an oblique incision closed with interrupted 5/0 Prolene sutures. After two weeks, mice were culled, and scaffolds were harvested and photographed for qualitative assessment.

### Cell Culture and Cytotoxicity Assay

Scaffolds were sterilised under ultraviolet light for 10 minutes in a tissue culture hood and placed individually into 96-well plates. Reset Vascular Endothelial Cells (R-VECs) were maintained in Endothelial Basal Medium (EBM; Lonza) supplemented with 2% foetal bovine serum (FBS) and EGMTM-2 SingleQuotTM growth factors (Lonza). Scaffolds and controls were seeded with R-VECs at densities of 5 × 10^3^, 1 × 10^4^, and 2 × 10^4^ cells/cm^2^ to determine the optimal cell seeding density for pre-vascularisation. Immediate and 72-hour viability and metabolic activity were measured using the MTS assay. MTS reagents were added to the culture medium and incubated for 2 hours, with absorbance read at 570 nm using an ELISA plate reader (Bio-Rad Gel DocTM EZ Imager).

### Embedding organoids into a pre-vascularised scaffold

A proof-of-concept experiment was conducted to embed blood vessel organoids alongside Reset Vascular Endothelial Cells (R-VECs) into pre-vascularised scaffolds, aiming to assess the effect of co-embedding on network density and stability. This was achieved using a simple suction method at a time point when both R-VEC networks were well established (day 5) and blood vessel organoids were in their angiosprouting phase (day 6). Culturing methods for both R-VECs and organoids are described elsewhere (Wimmer et al. 2019, Palikuqi et al. 2020).

R-VECs at a density of 1 × 10^4^ cells/cm^2^ were mixed with ten or more organoids in a total volume of 1.5 mL of organoid culture medium, which provided the necessary growth factors for R-VEC nutrition. One to two scaffold discs were placed into a 5 mL syringe by removing and then replacing the plunger. A syringe cap was attached to the syringe tip, and the cell-organoid suspension was introduced into the syringe from the base.

The plunger was carefully reinserted, generating positive pressure. The syringe was turned upright, and the cap was removed to allow the plunger to be pushed to the 1 mL mark without fluid loss. The syringe cap was then replaced. Aspiration and plunging were performed 4–5 times to facilitate the infiltration of media and organoids into the scaffold.

This vacuum-pressure method effectively mounted cells and tissue constructs into the internal spaces of the scaffold using standard syringe equipment. Care was taken to avoid inadvertently expelling media from the tip during plunger insertion and to remove excess air by briefly removing and replacing the cap. Alternatively, a three-way stopcock could be used, although it introduces greater dead space that consumes more cells. A 5 mL syringe was selected for its moderate vacuum and plunge pressures, as well as sufficient space for the scaffolds. Smaller syringes produce lower vacuum but higher positive pressures, which could damage cells and tissues.

### Statistical analysis

A minimum of 3 scaffolds were analysed per parameter variant. Data is presented as mean

+/-standard error of the mean (SEM). One-way ANOVA was used to analyse differences between samples using the Graphpad Prism 9 software. Differences were considered statistically significant when p-values <0.05.

## RESULTS

### Vessel density

Surface vascularity mapping was conducted to evaluate the vascularisation of various scaffold compositions and porosities (Figure 3). The average surface vascularity percentages for each scaffold type are summarised in Figure 4.

**FIGURE 1:**
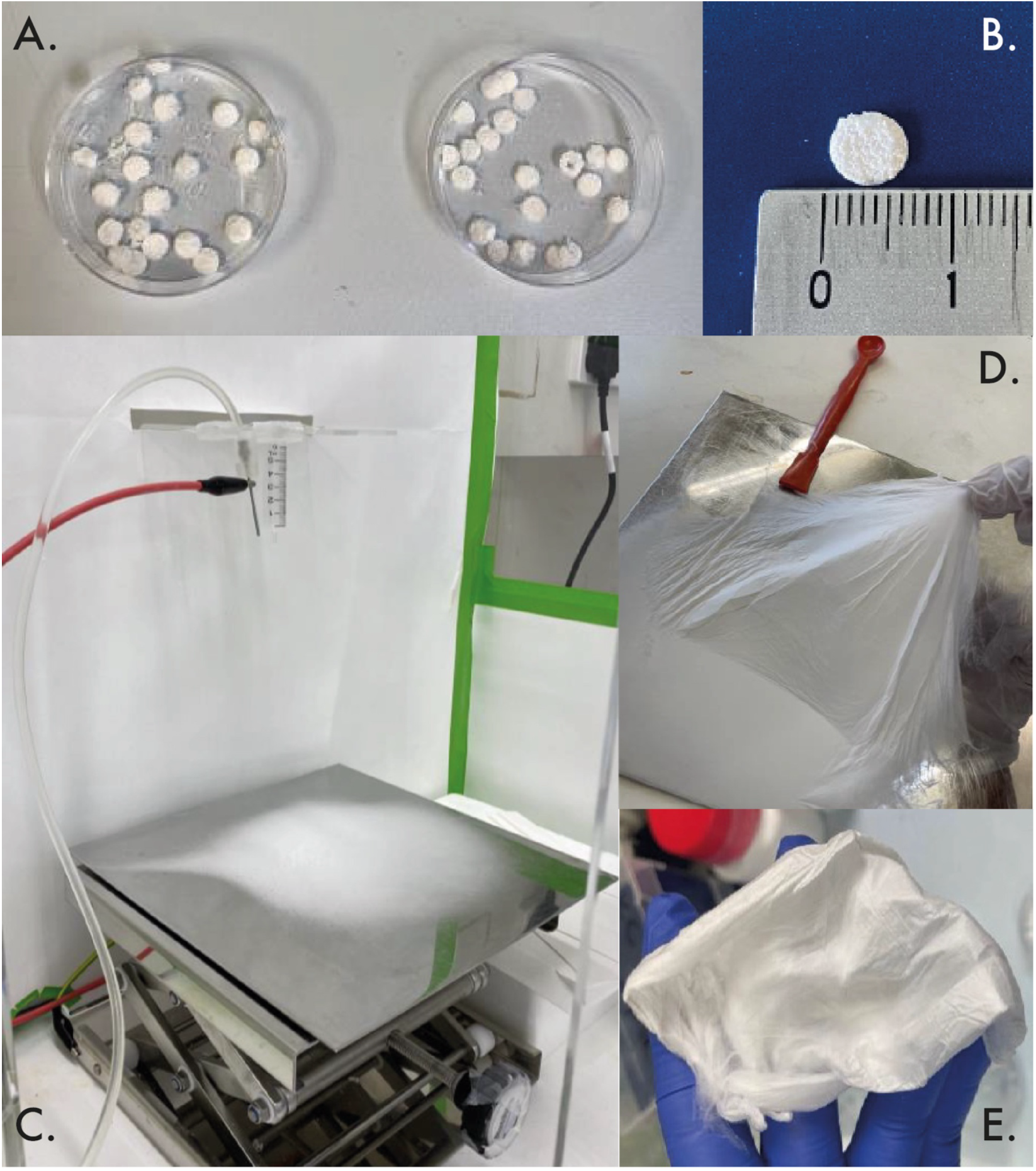
Fabrication process to create shapeable-PLGA or PLGA-collagen scaffolds. A,B. Size and shape of vacuum-dried discs produced using porogen-leaching. C. Set-up for electrospinning with suspended nozzle above collection plate. D, E. Retrieval of spun sheets that can be cut/shaped.

**FIGURE 2:**
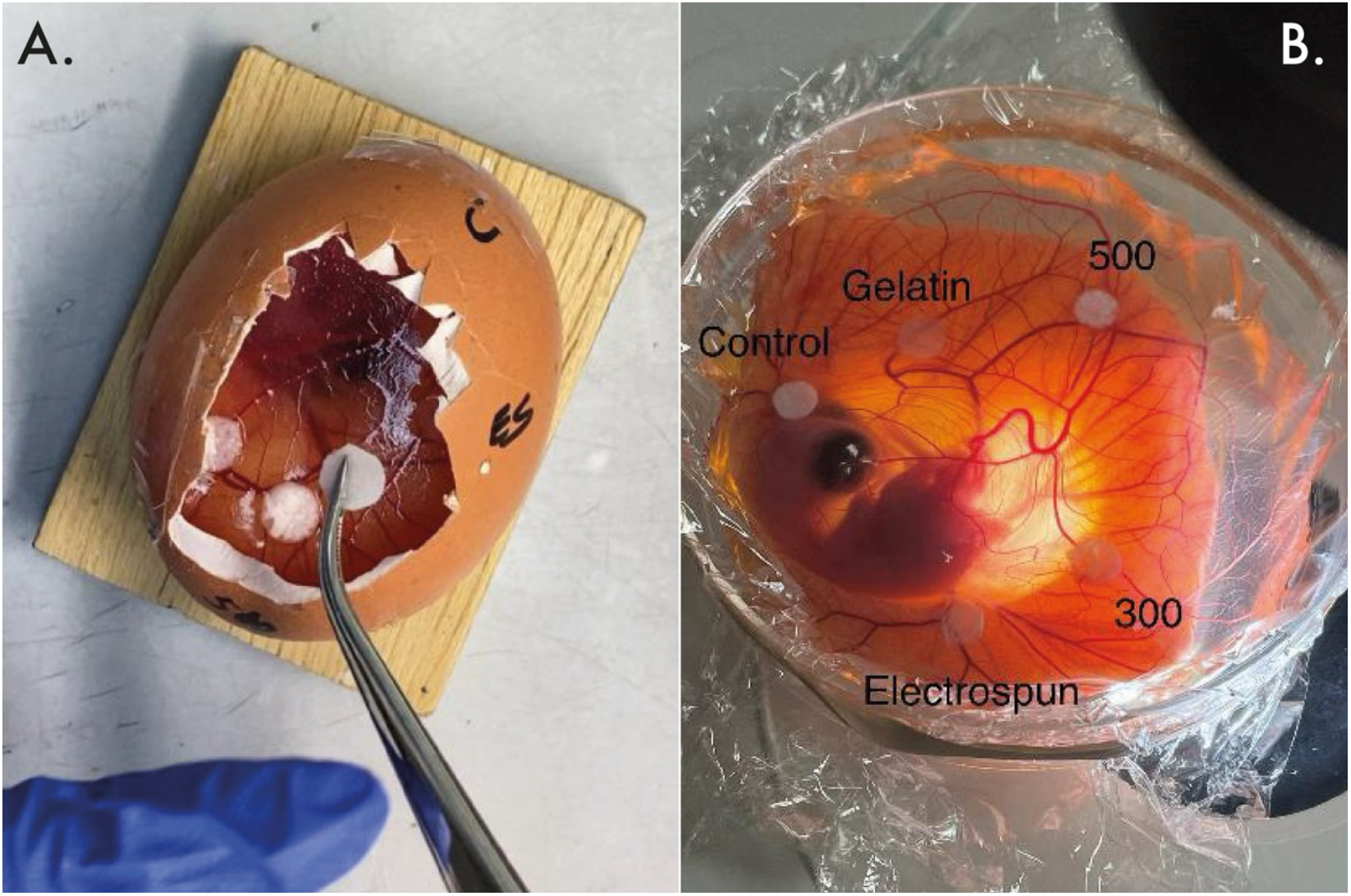
CAM-mounting technique. A. In-ovo method where scaffold discs are placed on CAM surface through a shell window. B. Ex-ovo method where CAM is released from shell and sits floating on a clingfilm surface within a jar.

**FIGURE 3:**
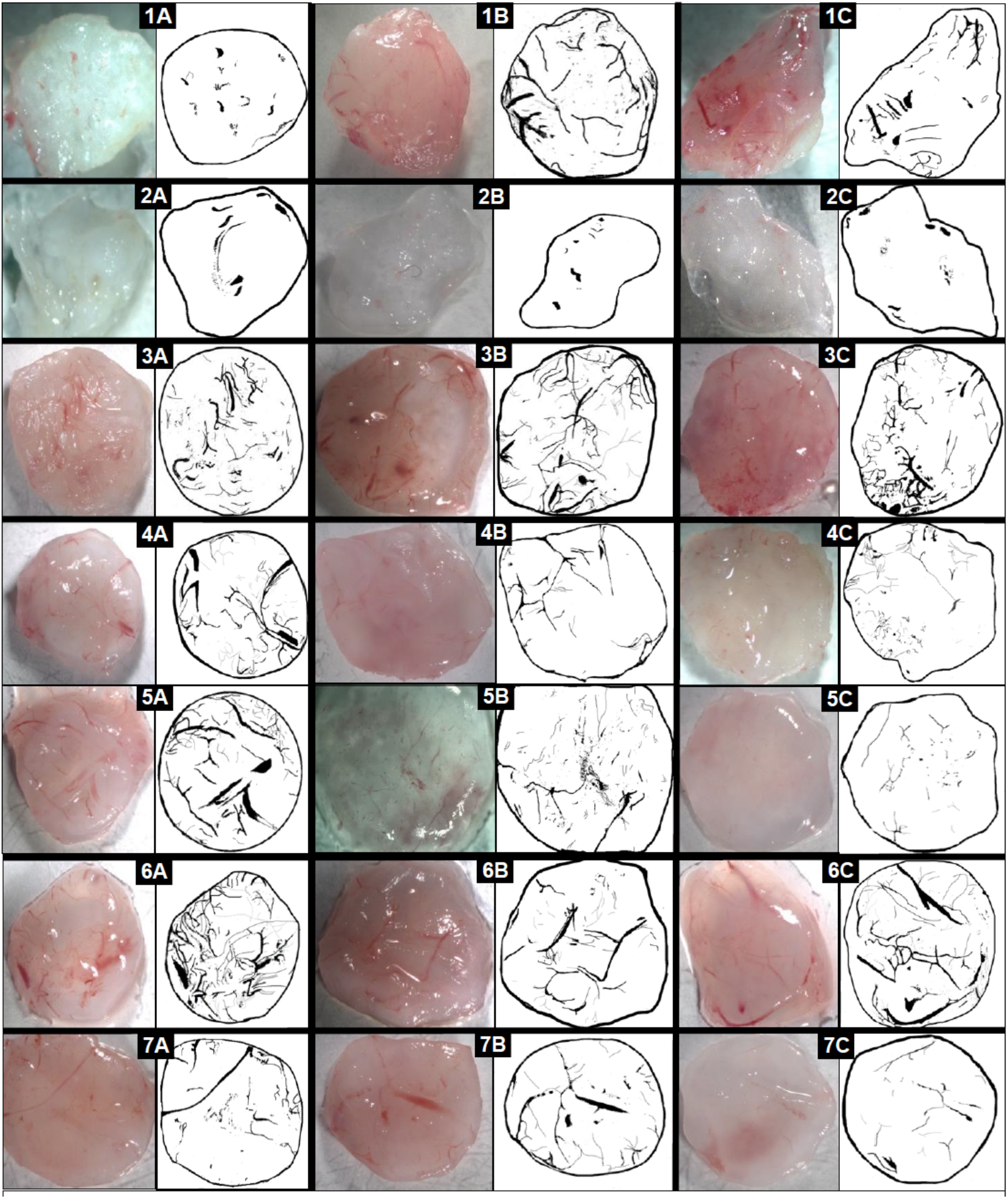
Vascular density mapping of scaffolds. Triplicate repeats of each scaffold vascular percentage generated. **1A-1C**. PLGA 75:25 with 300um porosity: 6.68%, 14.45%, 6.20% respectively. **2A-2C**. 75:25 with 500um porosity: 2.87%, 1.57%, 1.89%. 82:18 with 300um porosity: 10.19%, 9.32%, 14.30%. **3A-3C**. 82:18 with 500um porosity: 13.38%, 6.68%, 6.97%. **4A-4C**. 82:18 with 250 and 500um porosity: 15.21%, 8.30%, 1.15% respectively. **5A-5C**. Electrospun 82:18: 17.29%, 11.45%, 12.58%. **6A-6C**. Control scaffolds of filter paper: 10.13%, 8.28%, 8.45%.

**FIGURE 4.**
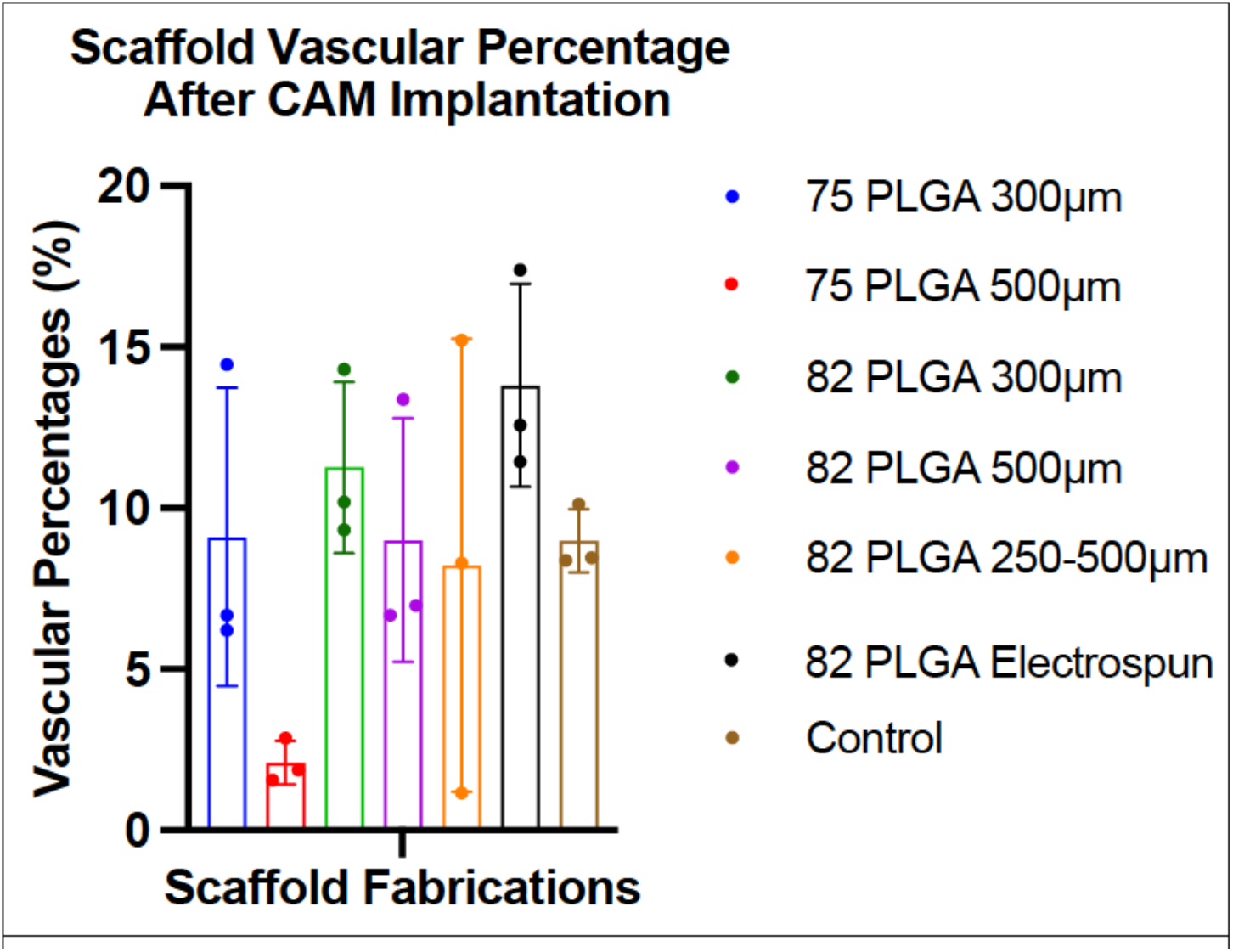
Histogram of vascular percentages. Data shows each of triplicate scaffold values from vascular mapping analysis. Data is mean + SD.

The electrospun PLGA 82:18 scaffolds exhibited the highest surface vascularity with an average of 14%. Among the porogen-leached scaffolds, PLGA 82:18 with 300 µm porosity showed a surface vascularity of 12%, while the 500 µm porosity variant had 9%. The PLGA 75:25 scaffolds demonstrated lower vascularisation, with the 300 µm porosity averaging 7% and the 500 µm porosity averaging 2.5%. Scaffolds with mixed porosity (PLGA 82:18 with 250 µm and 500 µm) had a surface vascularity of 8%, comparable to the control litmus paper, which also averaged 8%.

These findings indicate that both the polymer composition and the fabrication method significantly influence surface vascularity. PLGA 82:18 scaffolds consistently supported higher vascular densities than PLGA 75:25. Electrospun scaffolds showed the greatest vascular ingrowth, which may be attributed to their fibrous, matrix-like architecture.

### MicroCT vascular penetration scanning and in-vitro implantation studies

Using MicroFIL perfusion techniques, we successfully visualised vascular penetration within the scaffolds via micro-computed tomography (Micro-CT) imaging. The radio-opaque contrast agent allowed for clear segmentation of blood vessels from the lower-contrast PLGA scaffold walls. MicroCT imaging confirmed robust neo-angiogenesis extending through the full scaffold diameter, demonstrating the integration of vascular networks into scaffold cores. As shown in Figure 5, the Micro-CT images illustrate the extent of vascular infiltration.

**FIGURE 5:**
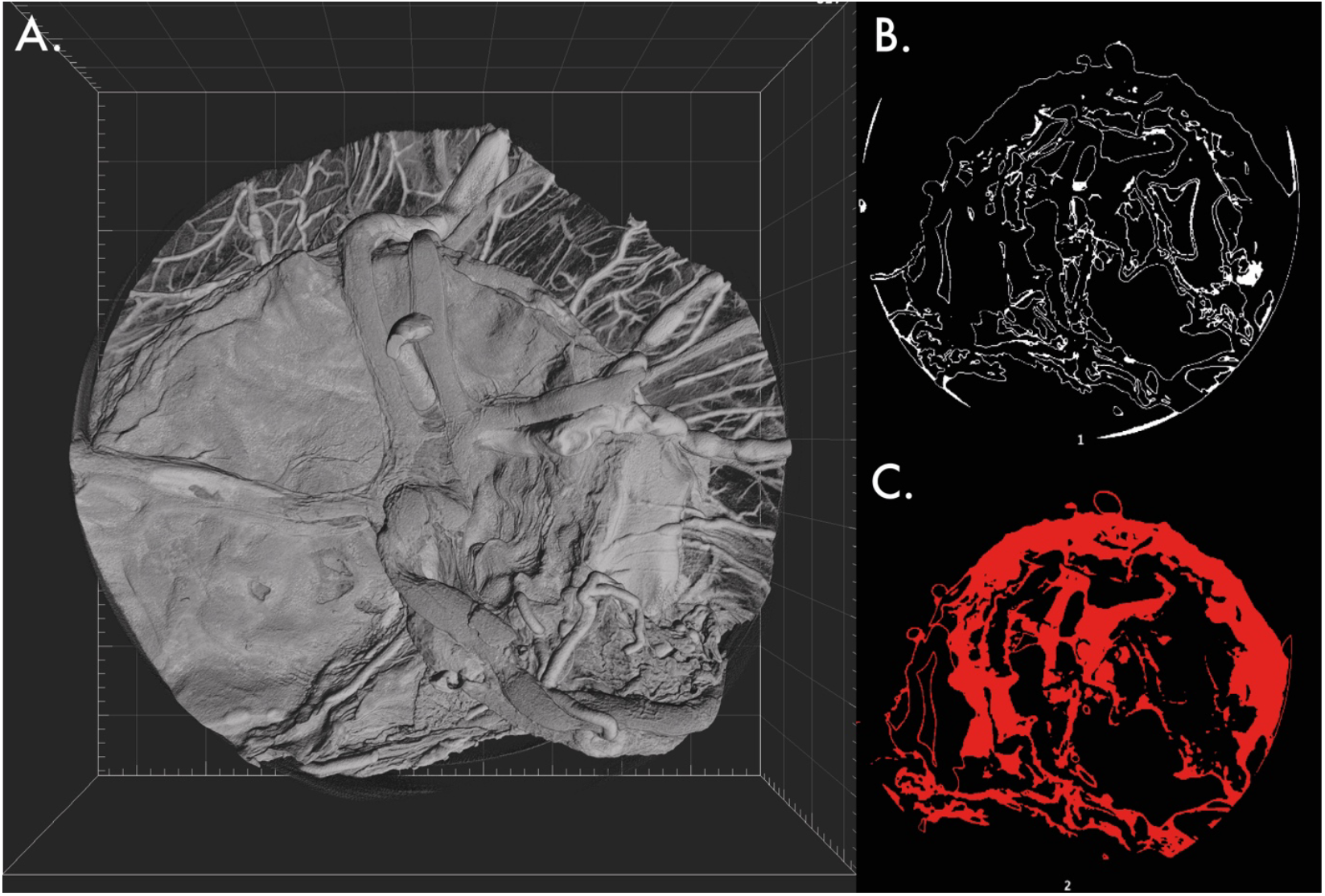
MicroCT images of MicroFIL-perfused scaffolds mounted on CAM assay. A. Topographical top-down image of the surface of the scaffold. Multiple small CAM-surface vessels can be seen entering the uppermost edge of the disc with larger calibre vessels grown over the top. B. Segmental image of the shape of internal scaffold walls. C. Segmental image of blood vessels within the scaffold.

**FIGURE 6:**
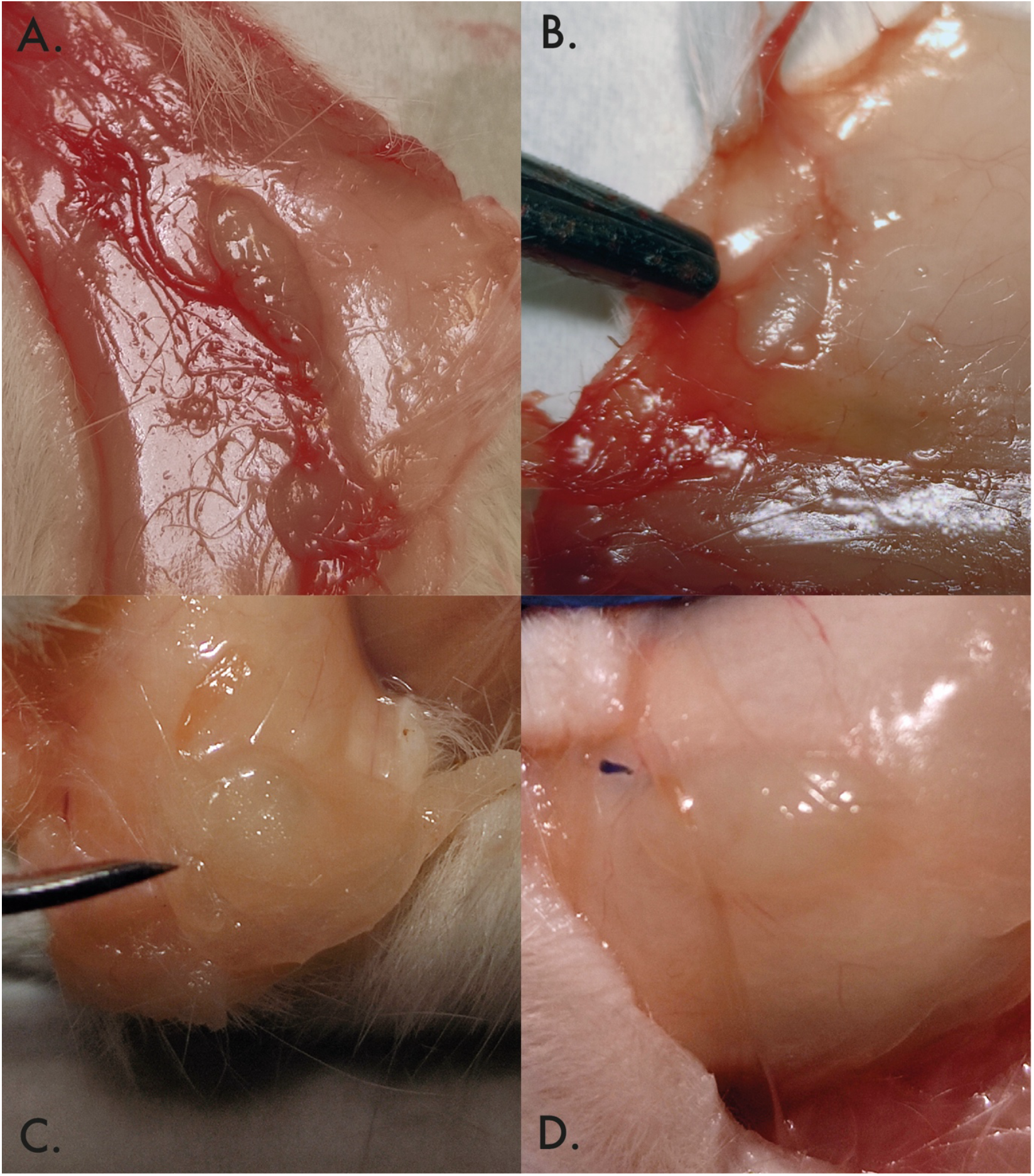
Photography of implanted bare PLGA scaffolds at 75:25 and 82:18 L to G composition ratios. Images show the change over four weeks to scaffold morphology. All scaffolds lost their starting disc-like morphology, moving towards a ‘rice-grain’ shape that may reflect mechanical pressure in the mouse dorsal skinfold alongside expected hydrolytic processes. **A**,**B**. 75:25 PLGA scaffolds were the most deformed by this process whilst the higher L ratio **C**,**D**. 82:18 PLGA scaffolds was more resistant to this morphology change. No scaffolds incited an inflammatory response, all being incorporated in a thin-layer of aponeurotic tissue with minimal surface vascularity. Data supports the principle of utilising higher L composition in PLGA scaffolds to maintain integrity both in culture phase and consequent implantation.

To assess the in vivo effects on scaffold morphology and degradation, bare PLGA scaffolds were implanted into the dorsal skinfolds of mice. Over a four-week period, all scaffolds exhibited a change from their initial disc-like morphology to a ‘rice-grain’ shape, which may reflect mechanical pressure within the mouse dorsal skinfold combined with hydrolytic degradation processes. No scaffolds incited an inflammatory response; all were incorporated within a thin layer of aponeurotic tissue with minimal surface vascularity. These findings support the utilisation of higher lactide composition (82:18 L:G ratio) in PLGA scaffolds to maintain structural integrity during both the culture phase and subsequent implantation.

### Seeding density

Optimal seeding density of R-VEC cells was determined to be between 10,000 to 20,000 cells/cm^2^, as these densities achieved the highest MTS absorbance readings both 2 hours after seeding (Day 1) and at 72 hours (Day 3). This trend was observed on both the electrospun PLGA 82:18 scaffolds and control filter paper. The results indicate robust cell attachment and metabolic activity at these densities, demonstrating the resilience and survival capacity of the R-VEC cell population under these conditions.

As shown in Figure 7, the 20,000 cells/cm^2^ density produced significantly higher absorbance values on Day 1 compared to the lower densities, particularly on the electrospun scaffolds. By Day 3, the absorbance values across the 10,000 and 20,000 cells/cm^2^ densities showed no significant difference, suggesting that the lower of these two densities is sufficient to sustain R-VEC metabolic activity over time.

**FIGURE 7:**
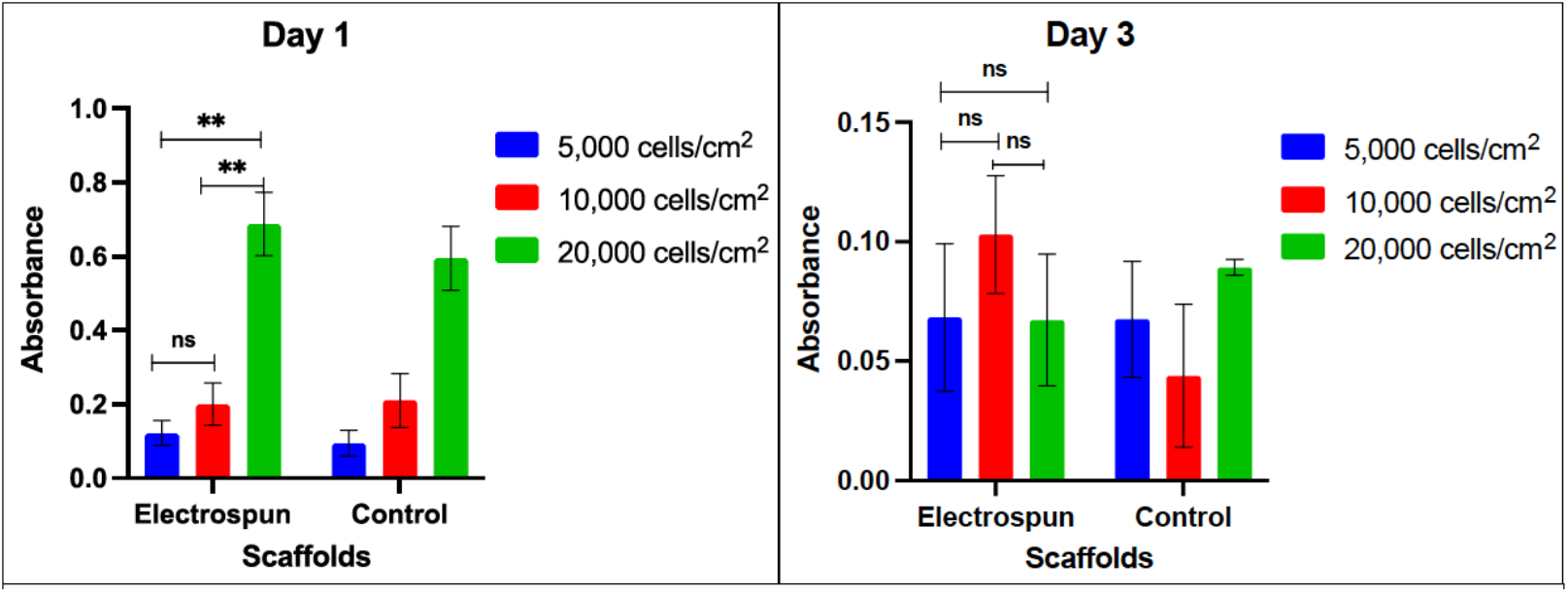
Histograms of cellular activity from MTS assay. R-VEC seeding was performed on both electrospun 82:18 scaffolds and control filter paper to determine the resilience of the cell type at different concentrations and at two time points. Absorbance was measured 2 hours after seeding (day 1) an at 72 hours after seeding (day 72). Data shows mean absorbance from 12 repeated samples for each seeding density and scaffold combination. Error bars represent standard deviation.

**FIGURE 8:**
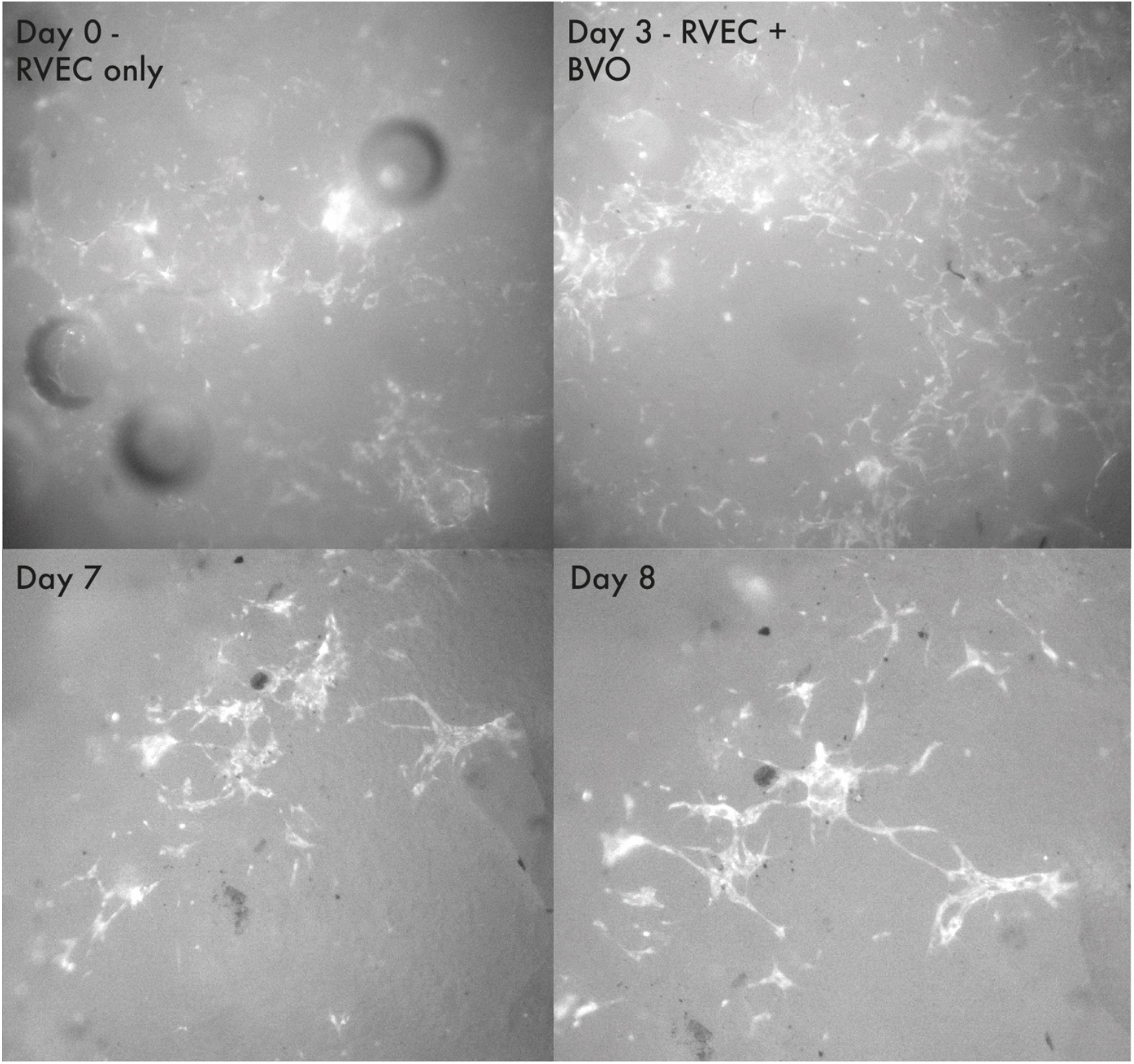
Embedding organoids onto a pre-vascularised scaffold. As proof of concept, RVECs were loaded onto PLGA 82:18 scaffold discs through submersion seeding (day 0). On day 1, scaffolds were retrieved and blood vessel organoids (BVO) were embedded through vacuum embedding and returned to culture wells. Vascular networks reached peak maturity on day 3, with gradual disintegration over the subsequent days. Images in the bottom row show appearance of sustained vascular networks around organoids (dark circles) at several places at day 7 and 8.

These results demonstrate the efficacy of electrospun scaffolds in supporting endothelial cell growth and highlight their potential as a platform for pre-vascularisation, particularly at seeding densities of 10,000–20,000 cells/cm^2^.

### Co-embedding organoids in a pre-vascularised scaffold

In a proof-of-concept experiment, pre-vascularised scaffolds seeded with R-VECs (20,000 cells/cm^2^) were co-embedded with blood vessel organoids (BVOs) to evaluate the impact of pre-vascularisation on organoid survival and vascular network stability. R-VECs formed robust vascular networks by Day 3, which gradually disintegrated between Days 5–7. When scaffolds were co-embedded with 20–30 BVOs, these networks remained visible around the organoids, suggesting sustained interaction between the vascular cells and organoids. This experiment demonstrates the feasibility of combining pre-vascularised scaffolds with organoids, providing a platform for future studies on vascular integration and organoid survival.

## DISCUSSION

Recent advancements in regenerative medicine and stem cell technologies have propelled interest in developing sophisticated tissue constructs, such as organoids and spheroids, that mimic the complexity of whole organs. Organoids offer transformative potential for disease modelling, drug discovery, and regenerative therapies. Yet, their success hinges on overcoming two key challenges: developing robust vascular networks to sustain constructs in vitro and ensuring rapid vascular integration (inosculation) upon implantation.. Current lab-on-chip platforms offer elegant but expensive solutions for studying vascularisation, yet their two-dimensional nature and lack of implantability limit their broader application. This study provides a scalable, low-cost approach to fabricating porous polymer scaffolds with demonstrated angiogenic potential. By leveraging R-VECs as a pre-vascularising cell type, we offer a practical solution for supporting organoid integration and survival.

The CAM assay served as a rapid, accessible platform to evaluate scaffold performance, enabling insights into angiogenesis and material biocompatibility. Importantly, the CAM model bypasses many of the ethical and logistical challenges associated with postnatal animal studies, while providing a robust orthotopic environment for testing (Ribatti 2012; Kohli et al., 2020). Across several scaffold compositions and porogen sizes, the higher 82:18 L:G ratio PLGA scaffolds consistently outperformed the lower 75:25 ratio scaffolds in maintaining their structural integrity, both on the CAM and in mouse dorsal skin-fold experiments. This durability is critical for future applications, as scaffolds must retain their form to support the sustained maturation and functionality of tissue constructs. The superior performance of 82:18 scaffolds underscores the importance of polymer composition in scaffold design for regenerative applications.

The R-VEC cell type proved highly effective for pre-vascularising these scaffolds, forming lumenised vascular networks capable of supporting advanced constructs. These cells, akin to embryonic vasculogenic endothelial cells, demonstrated resilience across seeding densities, with 20,000 cells/cm^2^ yielding optimal viability and metabolic activity in both short- and long-term assays. Furthermore, co-embedding R-VECs with organoids illustrated the scaffolds’ capacity to integrate with and stabilise vascular networks around organoids, supporting the construct’s survival over time. These findings highlight the potential for using pre-vascularised scaffolds as a platform for hosting complex tissue constructs prior to implantation (Palikuqi et al., 2020).

The integration of MicroFIL perfusion and micro-CT imaging provided a novel solution for evaluating vascular penetration within polymer scaffolds. Unlike conventional microscopy techniques, which struggle to visualise internal structures of polymeric materials, this approach leveraged the chick embryo’s natural perfusion pump to deliver contrast agent throughout the vascular network. The resulting imaging revealed deep internal vascularisation in R-VEC-seeded scaffolds, which contrasts with previous studies reporting only peripheral vessel inosculation (Woloszyk & Mitsiadis 2017). This advancement offers a powerful tool for assessing scaffold vascularity and construct viability, paving the way for improved preclinical evaluation of regenerative therapies.

This study integrates established fabrication, vascularisation, and imaging techniques into a scalable and cost-effective framework for producing pre-vascularised scaffolds. By combining R-VEC pre-vascularisation with scaffold porosity optimisation, we provide a practical solution for advancing organoid and tissue construct research.

## CONCLUSION

This study presents a robust workflow for developing pre-vascularised scaffolds capable of supporting advanced tissue constructs. By optimising scaffold porosity and polymer composition, we demonstrated that high L:G ratio (82:18) PLGA scaffolds maintain their structural integrity under in vivo conditions, making them suitable for regenerative applications. The R-VEC cell type, seeded at 20,000 cells/cm^2^, proved effective as a pre-vascularising population, forming self-organising vascular networks that enhance scaffold functionality and facilitate organoid integration.

Additionally, the combination of MicroFIL perfusion and micro-CT imaging provided an innovative method for evaluating scaffold vascularity, revealing deep internal vascular networks capable of supporting organoid survivability. These findings underscore the potential for scalable, cost-effective platforms that enable pre-vascularisation and organoid integration for translational research and regenerative therapies. Together, these approaches represent a significant step toward realising the potential of organoid-based regenerative medicine.

